# Improved Pathogenic Variant Localization using a Hierarchical Model of Sub-regional Intolerance

**DOI:** 10.1101/431536

**Authors:** Tristan J. Hayeck, Nicholas Stong, Charles J. Wolock, Brett Copeland, Sitharthan Kamalakaran, David Goldstein, Andrew Allen

## Abstract

Different parts of a gene can be of differential importance to development and health. This regional heterogeneity is also apparent in the distribution of disease mutations which often cluster in particular regions of disease genes. The ability to precisely estimate functionally important sub-regions of genes will be key in correctly deciphering relationships between genetic variation and disease. Previous methods have had some success using standing human variation to characterize this variability in importance by measuring sub-regional intolerance, i.e., the depletion in functional variation from expectation within a given region of a gene. However, the ability to precisely estimate local intolerance was restricted by the fact that only information within a given sub-region is used, leading to instability in local estimates, especially for small regions. We show that borrowing information across regions using a Bayesian hierarchical model, stabilizes estimates, leading to lower variability and improved predictive utility. Specifically, our approach more effectively identifies regions enriched for ClinVar pathogenic variants. We also identify significant correlations between sub-region intolerance and the distribution of pathogenic variation in disease genes, with AUCs for classifying *de novo* missense variants in Online Mendelian Inheritance in Man (OMIM) genes of up to 0.86 using exonic sub-regions and 0.91 using sub-regions defined by protein domains. This result immediately suggests that considering the intolerance of regions in which variants are found may improve diagnostic interpretation. We also illustrate the utility of integrating regional intolerance into gene-level disease association tests with a study of known disease genes for epileptic encephalopathy.

## Introduction

Accurate identification of pathogenic mutations is a central challenge in medical genetics and key to establishing accurate genotype-phenotype associations. Such an accurate classification requires one to consider not only the characteristics of the variants themselves, but also the genomic context in which they are found. Not all parts of a gene are of equal importance to viability and health. A variant that severely disrupts the function of one part of a gene may have a very different impact from a variant that severely disrupts another part of the same gene. This fact is also born out when one looks at distributions of disease mutations, which often cluster in specific sub-regions of genes.^1,2^ While variant level measures of functional impact are useful, their value can be increased by integration with information about the surrounding sequence context..

Cross species conservation^3^ has often been used to quantify sequence importance. A genomic region that is strongly conserved across species is likely subject to purifying selection, and therefore more likely to result in disease when disrupted. However, the conservation approach breaks down when the functionality of the sequence is human specific. To overcome this, Petrovski et al^4^ advanced an empirical measure of purifying selection in the human lineage. Using standing variation in the human population, they quantified the “intolerance” of genes by the observed depletion of common functional variation relative to expectation given the total amount of variation in the gene. Others have advanced similar gene-level scores, including “constraint”^5^ and “vulnerability”^6^ scores. These scores have proven useful in prioritizing genes by likelihood of being disease-related. However, they do not aid interpreting regions within genes, since they only score the entire gene and not its sub-units.

Sub-regional versions of intolerance and constraint scores have been proposed.^3–5,7,8^ Sub-RVIS,^7^ for example, partitions genes into functional subunits (e.g., exons or domains), and then measures the depletion of common functional variation, relative to expectation, within them. However, a limitation of sub-RVIS is that it does not consider the fact that each sub-region lies within a gene and therefore has natural relationships to other sub-regions in the same gene. It is already difficult to detect functional depletion on a gene-wide level, so analyzing smaller regions of genes adds to the challenge.^9^ If a sub-region is small, or the amount of variation is low, estimates of intolerance are highly variable and often inaccurate.

Since domains and exons are nested within genes, it is natural to model the relationship between subunits using a hierarchical model. Here, we improve existing methods and develop a statistical intolerance framework that is able to jointly model genome-wide, genic, and sub-regional effects. The hierarchical model, a Localized Intolerance Model using Bayesian Regression (LIMBR), facilitates the borrowing of information across a gene, leading to stabilization of estimates, lower variability, and improved predictive utility. As a result of this approach LIMBR shows much less bias due sub-region size than previous methods. To evaluate the utility of our approach we show that the intolerance scores are significantly associated with whether regions carry pathogenic mutations and are highly predictive, in particular, of *de novo* missense variants in Online Mendelian Inheritance in Man (OMIM) disease genes. Finally, we illustrate the utility of integrating regional intolerance into gene-level disease association tests with a study of known disease genes for epileptic encephalopathy, where results suggest that considering the intolerance of regions in which variants are found may improve power for gene discovery.

## Materials and Methods

### Bayesian Hierarchical Model

We establish a Bayesian hierarchical model explicitly characterizing depletion in funtioanl variation at both the gene and sub-regional level. As with sub-RVIS, we estimate intolerance within a genic sub-region by the departure, within the region, from the expected number of missense variants given the total number of variants (synonymous and functional) within the sub-region. However, unlike sub-RVIS, we use a hierarchical model that explicitly nests sub-regions within genes to further decompose this residual variation into gene-level and sub-region components. Figure 1 depicts the approximate geometric interpretation of this approach. The hierarchical model allows information to be shared across sub-regions, stabilizing intolerance estimates.

**Figure 1.**
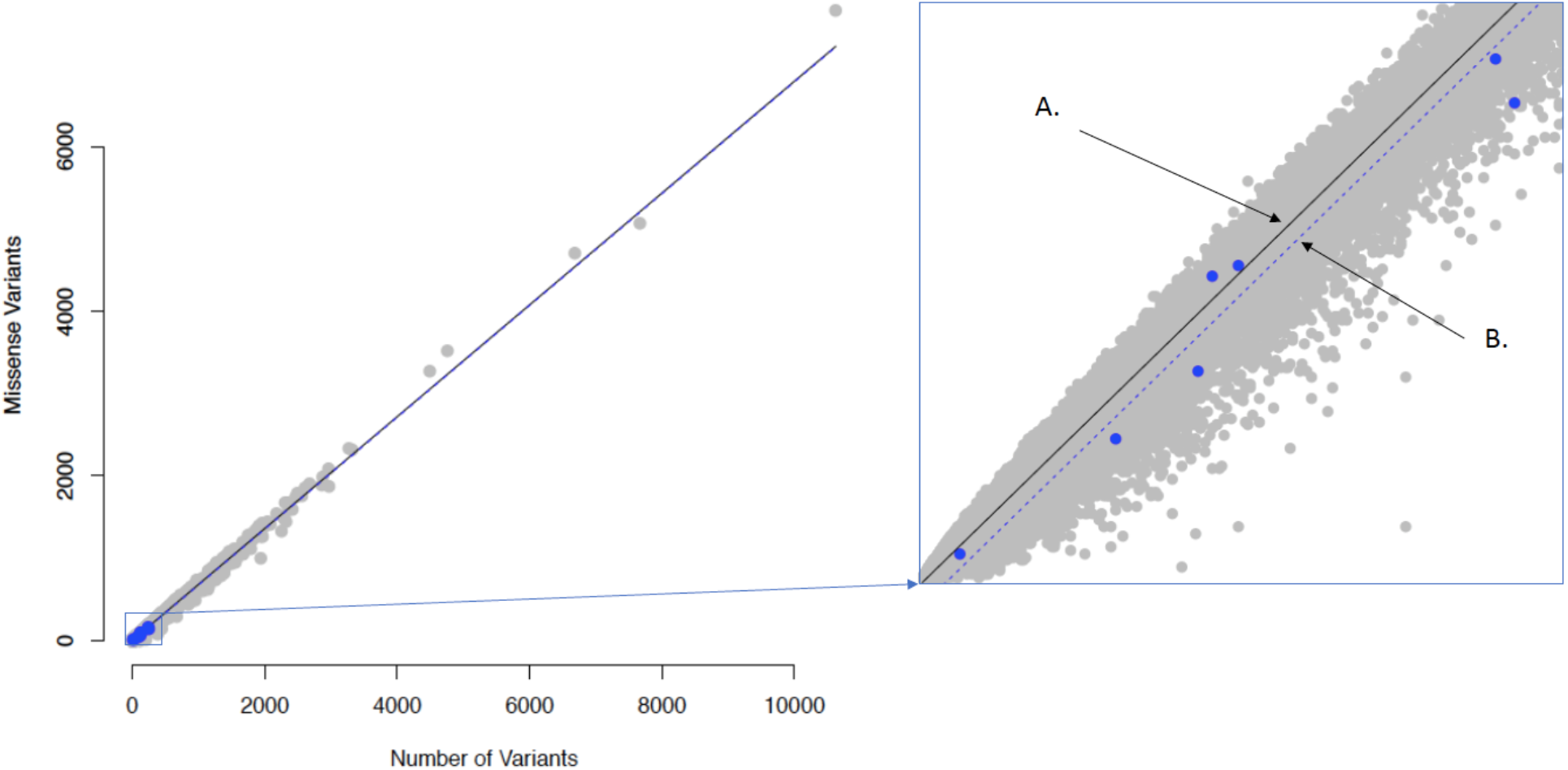
Relative missense variation versus total variation across domains. In blue are the SCN1A domains as an example while the grey dots correspond to all the other domains, where the genome wide average missense variation versus total variation is plotted as a black solid line (A). The offset average gene level trend for SCN1A is plotted as a blue dotted line (B) and can be seen more clearly in the exploded panel. Fitting a Bayesian hierarchical model allows for sharing of information across sub-regions, pulling the sub-region level terms towards the genic average.

We use a two-level linear mixed effects model with a single intercept for the genetic effect and two levels of random effects. Specifically, we model the number of missense variants within the *i*th sub-region of gene *j*, y_ij_, as a function of the total number of variants within the *i*th sub-region of gene *j,x_ij_*, through the following regression model, 
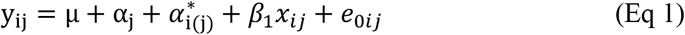
where µ is a genome-wide intercept, *α_j_* is the average deviation from µ for gene j, and *α*_i(j)_ is a deviation for the ith sub-region nested within gene j and 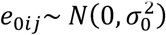.

We model *α_j_* and α_i(j)_ as random effects within a Bayesian framework. We assume standard normal priors for *α_j_* and α_i(j)_ but allow for separate prior variances for α_i(j)_ for each gene *j*. We begin by assuming an inverse gamma prior for the variances. Specifically, we choose *α_j_*∼*N*(0,σ^2^) with hyperparameters σ^2^∼*Inv-Gamma*(*ϵ,ϵ*) and *ϵ*∼*Uniform*(*δc*), where *δ* is a small positive constant and J is a large positive constant to induce a diffuse prior. For the sub-region parameters, we use a similar structure but with a separate variance for each gene, i.e., we choose 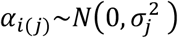 with hyper-parameters 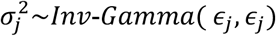 and *ϵ_j_*∼*Uniform*(*δ, c*). Note that by allowing for a gene-level variance, the *α_ij_*_(s)_ can be shrunken back to the gene level intolerance when there are no large differences between sub-region or when data is sparse. This will decrease the variability of the *α_i(j)_*_s_, leading to more stable intolerance estimation.

After initial testing the posteriors across different chains appeared to be volatile in certain settings, where the within versus across chain variance for some variables was above 1.1. To improve the ergodicity, *α_i_*(_*j*_) was set to zero for genes with 2 or fewer sub-regions, this is effectively just collapsing genes with only 2 sub-regions eliminating any inflated within versus across chain variance. Further, it is known that the hierarchical model can be augmented, sometimes referred to as noncentral^15–17^ or ancillary augmentation^18^, and similar methods are known to improve performance.^19,20^ So, we introduced an additional hyper parameter *v*∼*N*(0,1) for 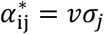to reduce autocorrelation.

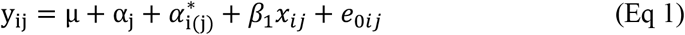

By introducing an auxiliary hyper parameter, the conditional variance structure is maintained while decoupling the random variables we wish to make inference on, 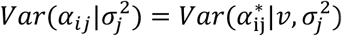. The final score that is used for the analysis is the posterior mode of the combined genic and sub-region terms.

The software was implemented in RStan^21^ interface of the Stan statistical modeling program, specifying the model in C++. Stan’s underlying software engine is a C++ program designed to be an efficient implementation of a Hamiltonian Monte Carlo (HMC) algorithm.^22^ HMC uses gradient information based on Hamiltonian dynamics, borrowing from physics, to better hone in on states of higher acceptance rates. Since the proposal step is leveraging more information it results in a faster convergence of the algorithm. A burn in of 1,000 with an additional 10,000 HMC steps across 5 chains was run for both domains and exons.

#### Assessing Prediction of Pathogenic Variants and Sensitivity to Sub-region Length

After calculating the scores, we perform assessment of the methods ability to predict pathogenic variants in several different scenarios. We directly compare the LIMBR scores against sub-RVIS to better understand if extending from a frequentist model, to the more involved and computationally intensive Bayesian model, improves prediction of pathogenic variance. It is also important to assess how sensitive the methods are to the relative sub-region lengths. For both sub-RVIS and LIMBR, the scores are regressed against the presence of ClinVar pathogenic mutations within each sub-region while controlling for mutation rate using logistics regression.

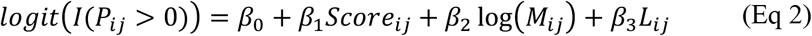

Where the sub-regions where considered pathogenic if they had at least one ClinVar pathogenic variant, *I*(*P_ij_* > 0). The log mutation rate, *M_ij_*, was included as a covariate and the model was fit both with and without sub-region length, *L_ij_*, as a covariate. The change in *β*_1_ was observed fitting the models with and without the sub-region length term in the model to see the potential effects of sub-region size.

Next, again using the calculated scores, a gene-by-gene assessment is run to test per-gene how well LIMBR predicts where within each gene pathogenic mutations are most likely to fall. For each gene the expected distribution of the pathogenic variants within the gene where calculated, based on the gene’s sub-regions’ mutation rates. First, regressing pathogenic variant counts within the sub-regions of genes against LIMBR scores and note the observed slope.

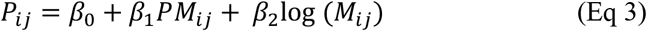

Regressing the model again but this time redistributing (permuting) the pathogenic variants into sub-regions by multinomial sampling, where variants are allocated to sub-regions proportional to the cumulative mutation rate of the sub-region, (see Methods for details). This effectively breaks any association between the location of the pathogenic mutations and intolerance. We repeat this process 1e4 times, characterizing the null distribution of the slope and thereby compute p-values by comparing the observed slope to permuted sets’ slopes.

Finally, we assess LIMBR’s utility in case-control studies to improve ability to detect gene-disease associations. The scores could be used as a weighting factor to increase power in disease related gene detection. A group-wise association test, similar to Madsen and Browning’s^24^ was constructed using the intolerance scores to weight the variants. An additive genetic model was used, where a genotype value of 0 corresponds to homozygous for the major allele, 1 to heterozygous, and 2 to homozygous for the minor allele. Assuming Hardy-Weinberg Equilibrium missing genotypes were imputed assigning each missing genotype a value of 2*p*, where *p* is the frequency of the minor allele in the cohort. For a gene with *n* exons, let *s_i_* be the intolerance score percentile of exon *i*. For all variants in exon *i*, the corresponding weight *w_i_* is given by: 
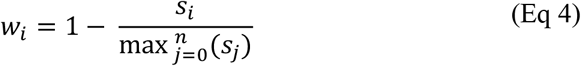

For the weighted model each individual’s genotype vector was multiplied by the weight vector and 1 for the unweighted model. A weight of 1 to all LoF variants since the intolerance score applies only to missense variants and LoF variants are expected to have the greatest relative effect on protein function.

The sum across each case individual’s weighted genotype vector yields an individual-level score, which is then summed to get the cohort wide score. Then 10 million permutations were generated of the score, where each time the case control status of each individual was permuted. The permutation p-value corresponds to the proportion of permutations that larger than the observed score with the true case control labels.

### Data and Quality control

The model is fit using 123,136 whole exome sequences from the genome Aggregation Database (gnomAD) across 17,765 genes.^10^ Loss of Function (LoF) variants are excluded from this analysis since LoFs can damage the resulting protein regardless of the sub-region in which they fall.^5,11,12^ Two sets of scores are calculated by fitting different definitions of sub-regions across genes, once with genic sub-regions defined by 172,123 exon boundaries and then again with sub-regions defined by 82,265 functional domains, both using the Conserved Domain Database (CDD).^7,13^ The filtered data from gnomAD first had to go through ‘PASS’ gnomAD criteria and further restricted to regions with at least 10x coverage in at least 70% of the samples. The model was regressed on missense variation versus all classes of functionality. Variants causing loss-of-function (LoF) damage the resulting protein, regardless of the sub-region, so we removed those predicted to be LoF. Variants that were annotated as either: splice acceptor variant, splice donor variant, stop gained, or stop lost were considered LoF. Indels were ignored. Exons were defined based on consensus coding sequence project release 20 (CCDS20).^25^ All coding regions of a gene were considered across all possible CCDS transcripts of a gene. Domains were defined in the Conserved Domain Database (CDD), and aligned to CCDS15.^7,25^ ClinVar^14^ pathogenic variants that were labeled as either pathogenic or likely pathogenic were used for the validation analysis and control variants were taken from 50,726 DiscovEHR^26^ samples, excluding variants also found in gnomAD and ClinVar.

## Results

### Exome Wide Predictive Ability of Intolerance Scores on Pathogenic Mutations

To better understand the advantage of the hierarchical approach, we examine the top and bottom 10% of the intolerance scores from both LIMBR and sub-RVIS and the number of bases spanned by these regions (Figure 2). For exon-level sub-RVIS, the average bases spanned for the top 10% intolerant exons is 452, compared to an average of 214 bases for the bottom 10% (Figure 2a: t_df=25397_ = 44.01 and p < 2.2e-16). For domains, the top intolerant domains average 1124 bases versus 559 bases for the most tolerant regions (Figure 2c: t_df=25397_ = 40.9 and p < 2.2e-16). Using LIMBR, the top 10% intolerant exons average 297 bases versus 286 bases for the most tolerant exons (Figure 2b: t_df=33322_ = 2.2, p-value = 0.034), and 717 bases versus 654 bases for domains (Figure 2d: t_df=16145_ = 4.4, p-value = 1.3e-5). LIMBR appears considerably less sensitive to sub-regional length; there is around a 10-base difference on average between the most and least intolerant exons, which is only marginally significant at an α = 0.05. In contrast, exon-level sub-RVIS has a highly significant difference of over 200 bases spanned between the most and least intolerant exons. As shown in separation of the sub-region length histograms for top 10% and bottom 10% scores points to a potential association between longer regions and higher intolerance scores in sub-RVIS.

**Figure 2.**
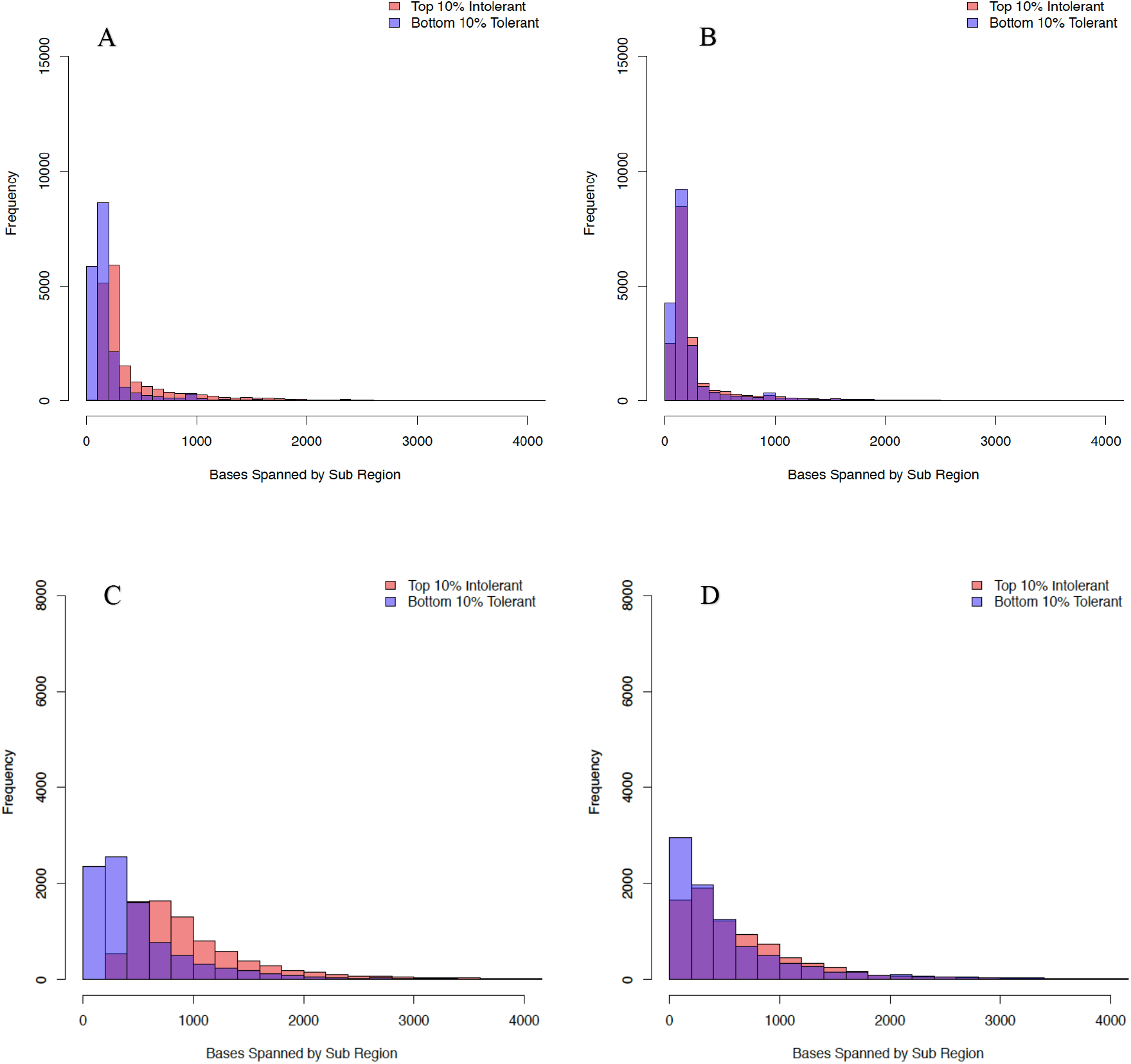
Comparing sub-RVIS versus LIMBR length distributions looking at the top and bottom 10% intolerant sub-regions. The histograms show the distribution of the number of bases spanned in sub-regions in top and bottom 10% of intolerance scores sets for sub-RVIS (exons A. and domains C.) and then the top and bottom intolerant LIMBR scores (exons B. and domains D.).

To explore whether LIMBR provides a benefit over sub-RVIS in terms of classification, we consider how well the resulting intolerance scores predict the presence of pathogenic mutations. Specifically, for both sub-RVIS and LIMBR, we regress the sub-region scores on the presence of ClinVar pathogenic mutations within these sub-regions while controlling for mutation rate using logistic regression (Eq 2). We see a significant association between exon-based intolerance scores and the presence of pathogenic variants for both LIMBR (p = 1.3e-29) and sub-RVIS (p = 2.8e-13), and less significant association for domains (LIMBR p = 1.6e-28, sub-RVIS p = 6.6e-67). Then we want to understand the scores ability to classify pathogenic regions in the context of region size and if there is some potential bias, as implied by Figure 2. To directly test for this, we fit a logistic regression model as described above, but with the addition of sub-region bases spanned as a covariate. An observed 49% (exons) and 11% (domains) changes in the estimate of sub-RVIS effect size after adding the sub-region length as a covariate, as opposed to around 14% (exons) and 2% (domains) changes for LIMBR. This indicates the possibility of strong influence of the size of the sub-region in the relative intolerance score using the sub-RVIS model.

We then consider how well a given intolerance threshold captures pathogenic mutations. Specifically, a given intolerance threshold defines both the total number of pathogenic mutations found in sub-regions as intolerant (or more) as the threshold as well as the total number of bases spanned by these same sub-regions. LIMBR appears to perform consistently better than sub-RVIS, capturing more pathogenic variants per base spanned (Figure 3). The analysis indicates similar results between exon-level LIMBR and domain-level LIMBR in terms of the number of pathogenic variants captured versus bases spanned (Figure 3).

**Figure 3.**
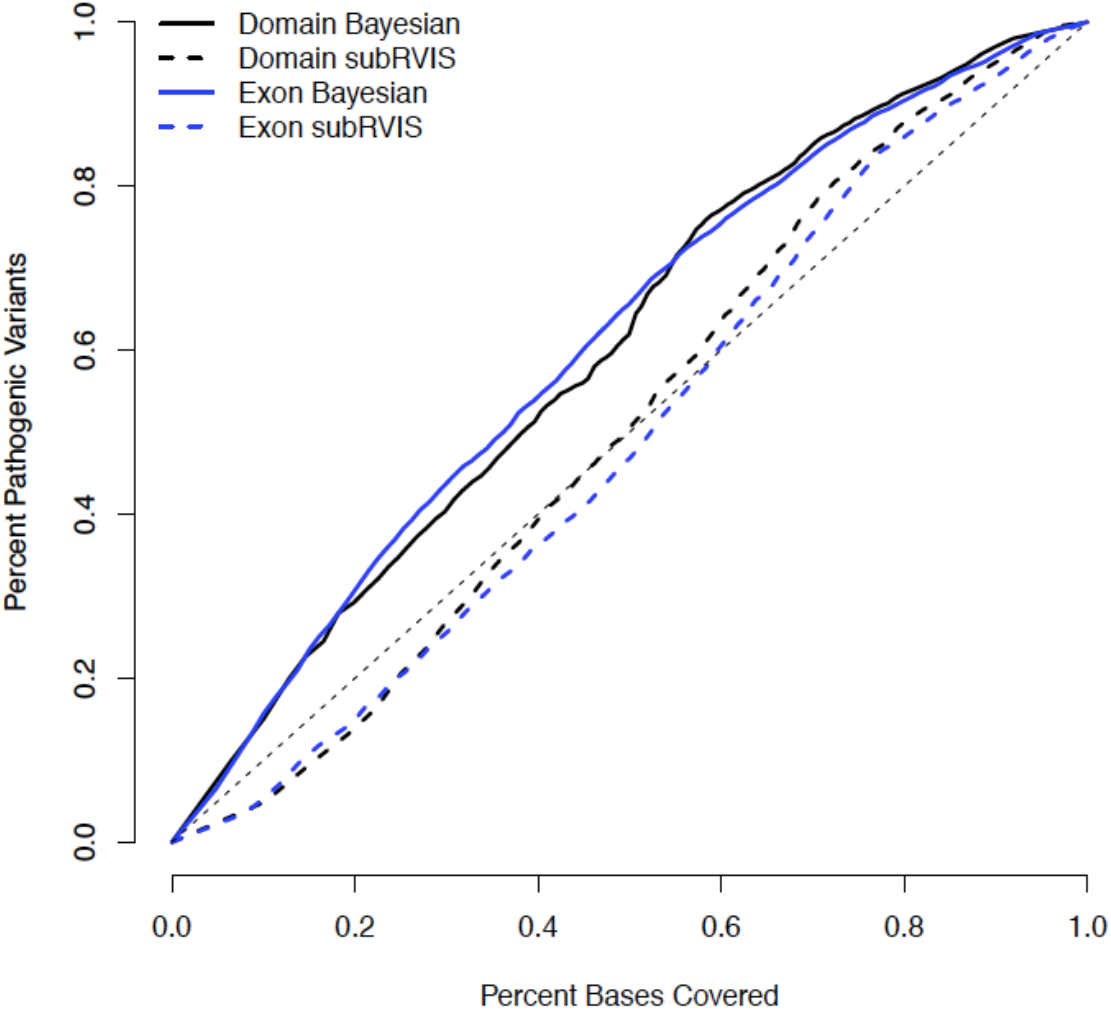
Comparing LIMBR and sub-RVIS ability to capture ClinVar pathogenic variants versus percent of bases spanned, restricting to OMIM genes. The genes are either broken up into exons (blue) or domains (black), thensub-regions are sorted based on either their sub-RVIS (dotted lines) or LIMBR scores(solid lines). At each percentile the percentage of ClinVar pathogenic variants captured relative to the percent of bases covered is compared between the two methods.

### Evaluation of impact of incorporating sub-regions using ClinVar Pathogenic Variants

We compare LIMBR to existing gene-level scores, including genic RVIS, pLI, and missense Z,^5,10,27^ beginning by focusing on OMIM genes. Since there is no explicit way of knowing if regions are benign we perform several comparisons against ClinVar pathogenic variants to help understand how well the methods classify variants. First, the percentile-sorted scores are used as thresholds to assess the ability of the methods to capture ClinVar pathogenic *de novo* missense variants versus benign sub-regions (Figure 4a-c for exons and Figure 4d-f for domains). In this analysis, LIMBR has the highest AUC (0.86) followed by missense Z (0.81), RVIS (0.78), pLI (0.76), and sub-RVIS (0.59) (Figure 4a). Similarly, for domains LIMBR has the highest AUC (0. 91) followed by sub-RVIS (0.81), missense Z (0.81), pLI (0.78), RVIS (0.76) (Figure 4d). LIMBR demonstrates at least a 5% and up to 10% improvement in AUC for classification of pathogenic missense *de novo* versus other methods. We observe similar results when using epilepsy and neurodevelopmental AD gene sets.

**Figure 4.**
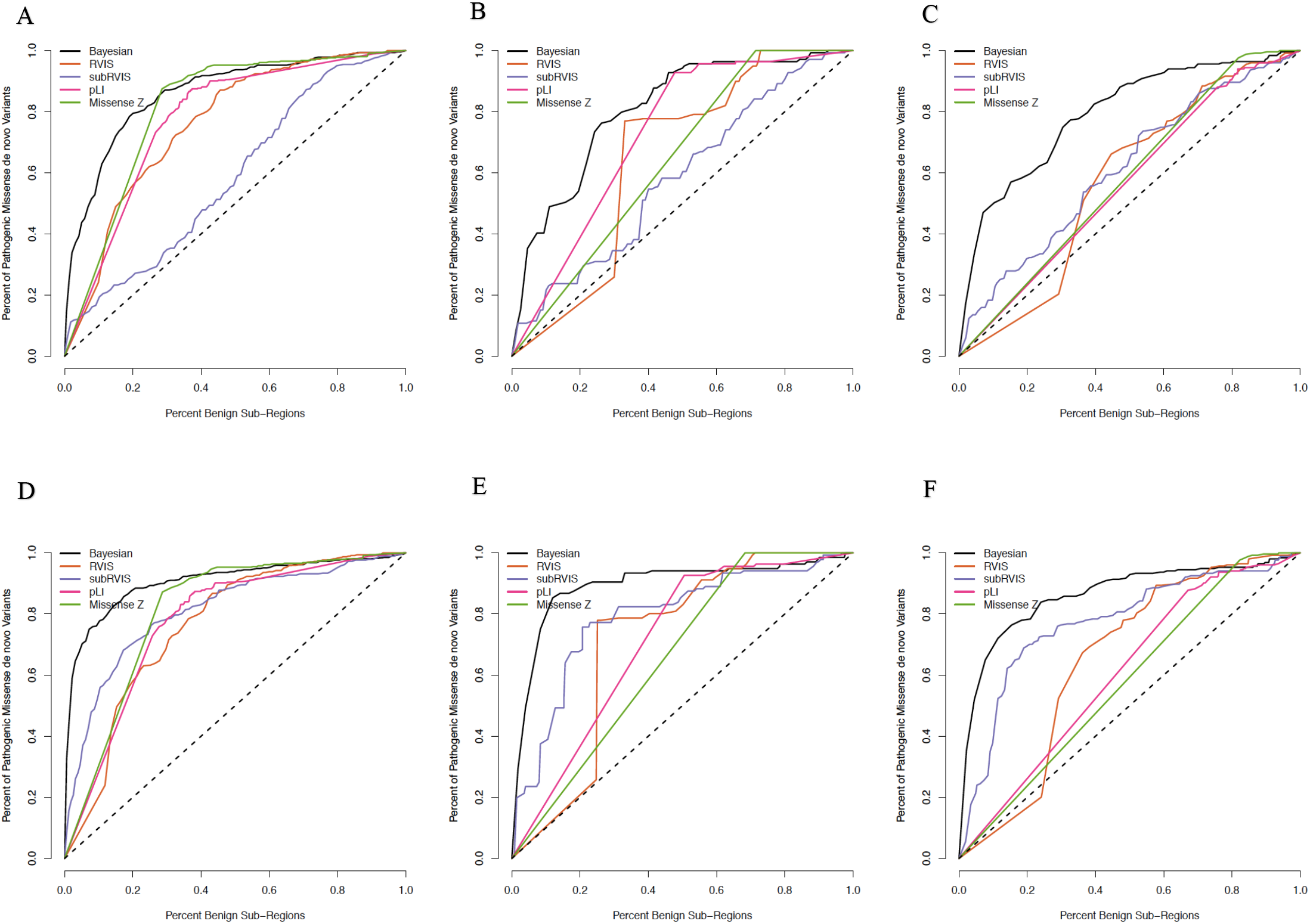
Performance of different methods’ ability to capture de novo pathogenicity across exons relative to benign variation. The methods are compared looking first at exons and the percent de novo missense variants versus benign exons restricting to a.) all OMIM genes c.) epilepsy gene set d.) neurodevelopmental autosomal dominant genes. Then for domain groupings, the methods are compared again with the percent de novo missense variants versus benign exons restricting to d.) all OMIM genes e.) epilepsy gene set f.) neurodevelopmental autosomal dominant genes.

Then no longer restricting to *de novo* we classify sub-regions as pathogenic or benign, where a sub-region is pathogenic if it contains at least one ClinVar missense pathogenic variant and benign otherwise. Scores are sorted by percentile and use it as a threshold to examine the percentage of pathogenic sub-regions versus benign sub-regions captured. We examine all OMIM genes, then restrict to either epilepsy genes or neurodevelopmental autosomal dominant (AD) genes (Figure 5a-c and Figure S1a-c for domains). For exons across all OMIM genes, LIMBR has the highest AUC (0.58) followed by sub-RVIS (0.56), missense Z (0.54), pLI (0.52), and RVIS (0.49) (Figure 5a) whereas for domains sub-RVIS performs best (0.62) followed by LIMBR (0.59), missense Z (0.56), RVIS (0.56) and pLI (0.53).

**Figure 5.**
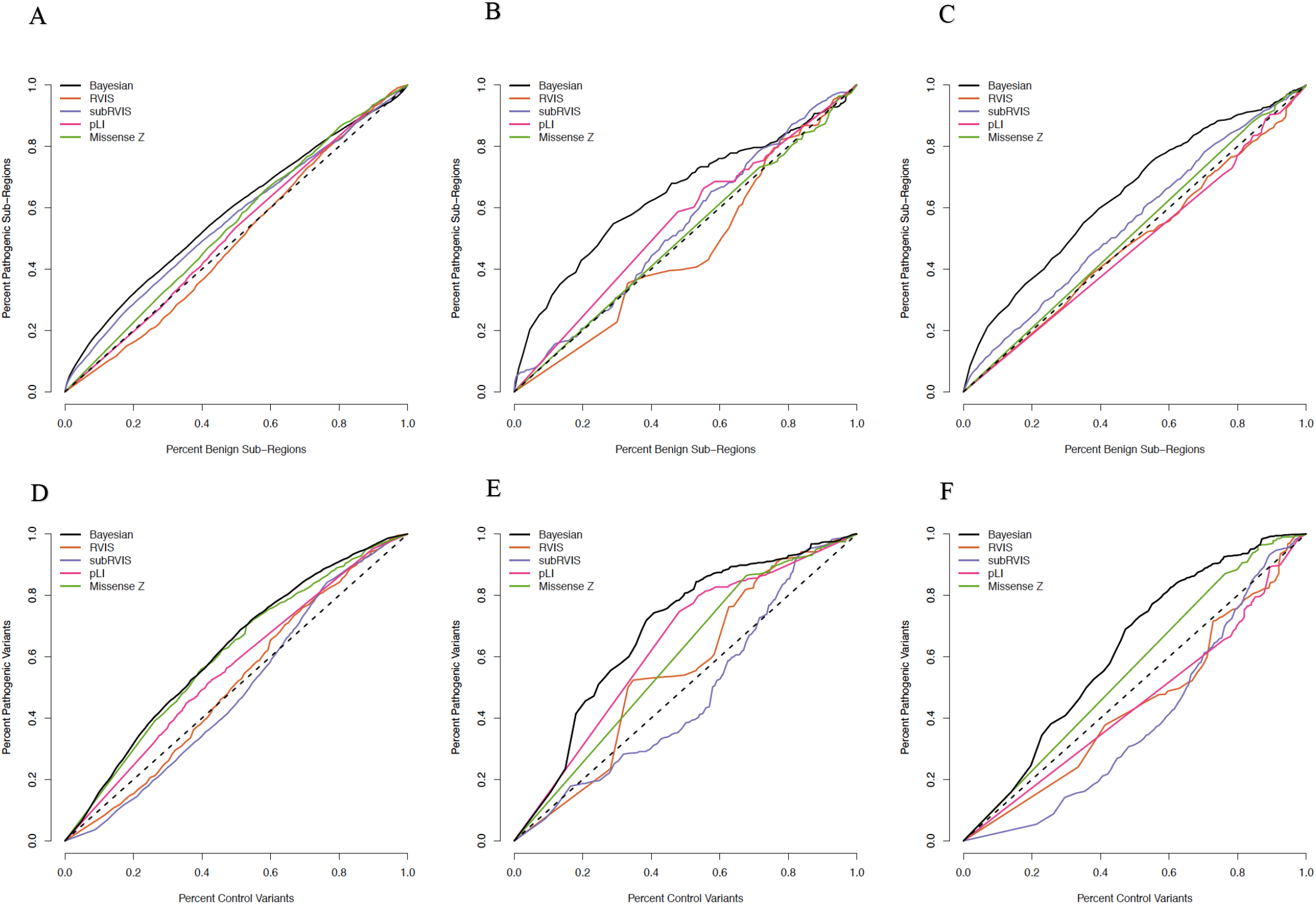
Performance of different methods’ ability to capture pathogenicity across exons relative to benign variation. LIMBR is compared against the other methods to see how well it classifies pathogenic exons versus benign exons restricting to a.) all OMIM b.) epilepsy gene set c.) neurodevelopmental autosomal dominant gene set. Then the percent pathogenic variants versus control variants captured by the different methods is compared restricting to d.) all OMIM e.) epilepsy gene set f.) neurodevelopmental autosomal dominant gene set.

Then the percentile-sorted scores are used as a threshold to examine the percentage of missense pathogenic ClinVar variants captured versus control variants, where control variants are defined as variants found in DiscovEHR^26^ control samples, excluding variants also found in gnomAD and ClinVar (Figure 5d-e and Figure S1d-e for domains). Despite sub-RVIS performing nearly as well as LIMR in terms of classifying sub-regions, when we examine the ability to classify pathogenic versus control variants, sub-RVIS performs the worst and LIMBR performs the best (Figure 5d and Figure S1d). We also considered the percent ClinVar pathogenic variants captured versus benign sub-regions (for exons Figure S2a-c and for domains Figure S31a-c) and the percent ClinVar pathogenic variants captured versus bases covered (for exonsFigure S2d-e and for domains Figure S3d-e). Similar results are observed when restricting to epilepsy genes (Figure 5b,Figure **5**e,S1b,S1e,S2b, and S2e) and neurodevelopmental AD genes (Figure 5c,Figure**5**f, S1c, S1f, S2c, and S2f), where LIMBR appears to demonstrate consistent improvement in capturing pathogenic variation. LIMBR results across different gene sets: epilepsy, neurodevelopmental, AD, neurodevelopmental AD, haploinsufficient, and recessive genes (Figure 6 for exons and Figure S4 for domains) are plotted for further comparisons of the gene groupings.

### Genes with Significant Association between Sub-Region Level Intolerance and Pathogenic Variant Placement

To test if the intolerance scores are significantly predictive of where pathogenic mutations fall within a given gene a permutation test is constructed. Using this permutation test (Eq 3), there are 259 genes that demonstrate significance at α = 0.05. Of the 259 significant genes, 41 are haploinsufficient, 76 are recessive, 101 are AD, 103 are neurodevelopmental, 41 are both neurodevelopmental and AD, and 15 are epilepsy genes (Table 1 and Figure 6). Examining gene sets in different organ groups from Database of Chromosomal Imbalance and Phenotype in Humans Using Ensembl Resources (DECIPHER)^28^, for face genes, 29 out of 252 demonstrate significant permutation tests (exact test p = 0.004), and for heart cardiovascular/lymphatic genes 32 out of 240 demonstrate significance (exact test p = 2.7e-4) (Table 1). This also includes 5 genes known to be associated with early-onset epilepsy: SCN1A (p-value <1e-5), SCN8A (p-value = 2e-4), CDKL5 (p-value = 0.011), PCDH19 (p = 4e-5), and KCNT1 (p-value = 0.006) (Figure 7).

**Figure 6.**
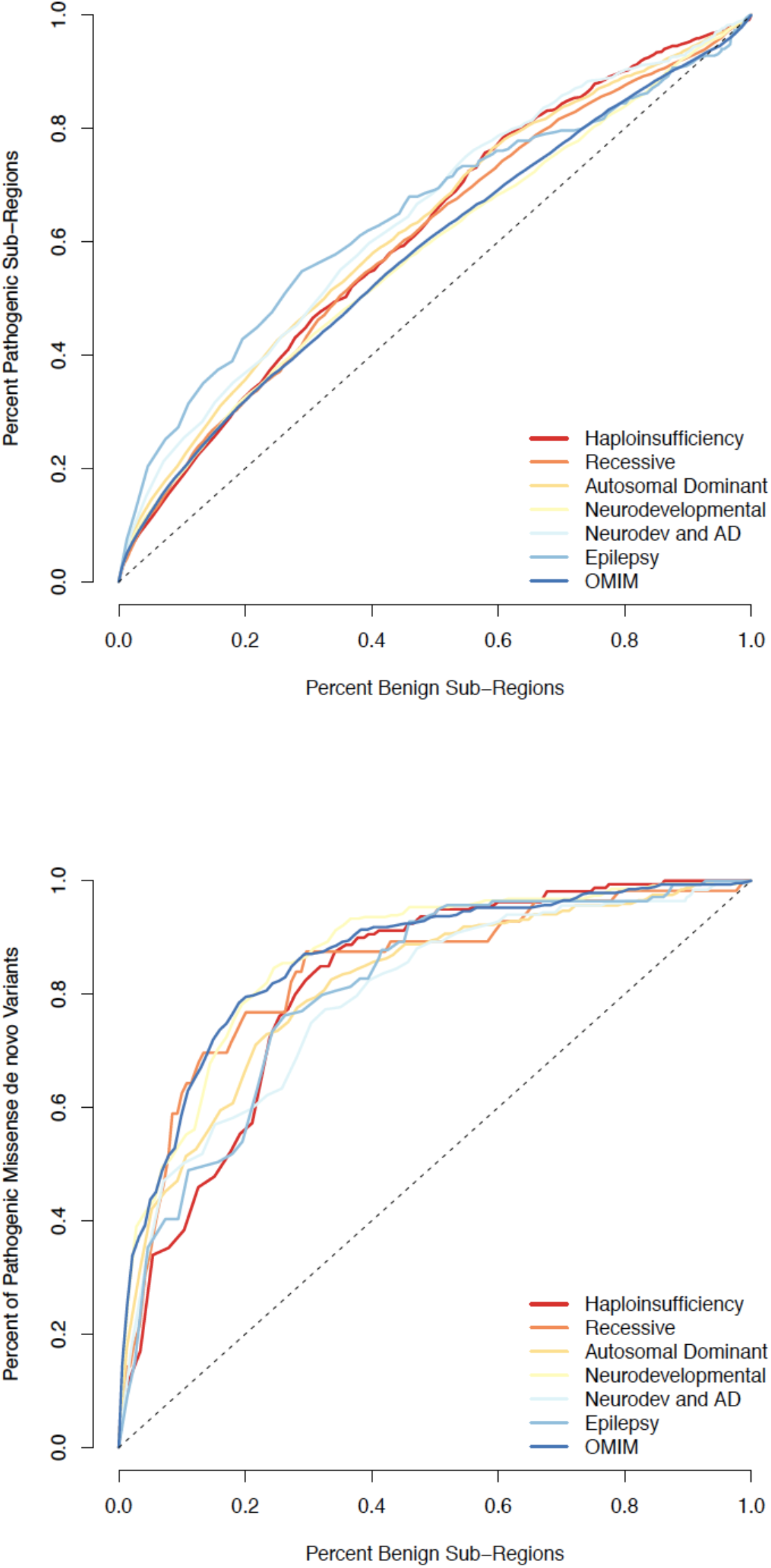
Performance of LIMBR across different gene sets. The LIMBR classification r a.) plotting exons with at least one pathogenic variant versus benign exons restricting to different OMIM genes sets (overlapping with set in Figure 4). Then similarly using to LIMBR percentile rankings of exons to see the b) percent pathogenic de novo variants relative to benign exons again restricting to different OMIM genes sets.

**Figure 7.**
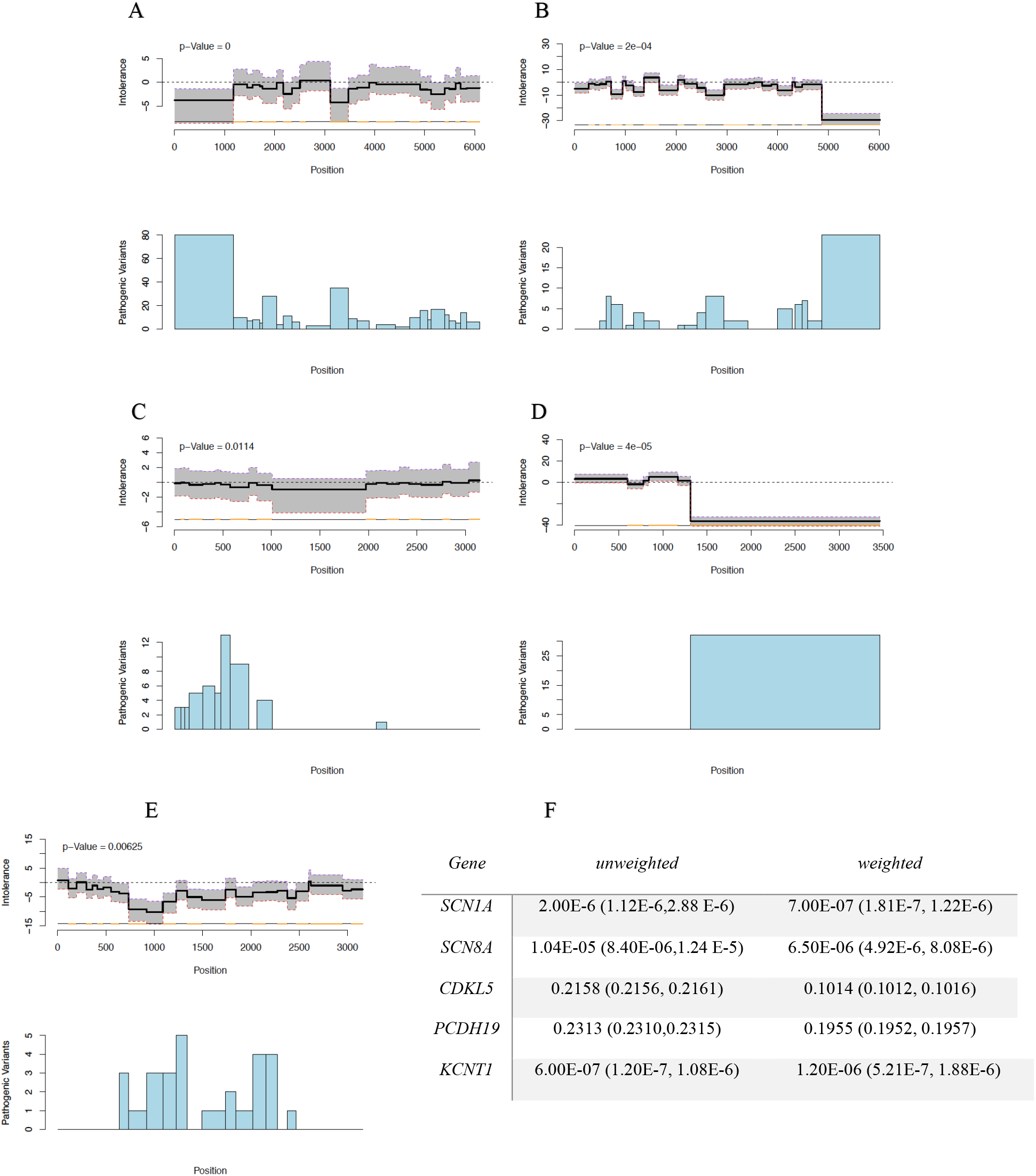
Localized genic intolerance to variation in key epileptic encephalopathy genes. The plots above are of the intolerance scores for, a.) SCN1A, b.) SC81A, c.) CDKL5 d.) PCDH19 and e.) KCNT1 with 95% credibility in grey across combined coding positions in all transcripts. The bar strip below that plot indicates when the start and end of a exon occurs. Below are the densities of ClinVar variants matched up at the corresponding genomic positions with the intolerance scores. Then f.) is a table that depicts a group-wise association test, both unweighted and weighted with the inverse percentile of the LIMBR intolerance, using a cohort of 488 epileptic encephalopathy cases and 12,151 unrelated controls from a previous rare variant collapsing analysis.

**Table 1.** Gene set enrichment of using gene level permutation test demonstrating significant association between exon level intolerance and pathogenic variant localization. The table above depicts different gene sets with the number of genes in each set, of those genes the number of them that are significant using the permutation test, and the exact test for observing that many significant genes in the sub-category relative to the total OMIM disease genes.

| *Gene Set* | *Number of*<br>*Genes In Set* | *Number Genes*<br>*Significant*<br>*Permutation Test* | *Exact*<br>*p-Value* |
| --- | --- | --- | --- |
| *Haploinsufficiency* | 175 | 41 | 2.04E-12* |
| *Recessive* | 817 | 76 | 0.027* |
| *Autosomal Dominant* | 1083 | 101 | 1.56E-11* |
| *Neurodevelopmental* | 1010 | 103 | 1.21E-07* |
| *Neurodevelopmental AD* | 252 | 41 | 2.08E-10* |
| *Epilepsy* | 100 | 15 | 2.66E-04* |
| *Bone Marrow/Immune* | 143 | 7 | 0.705 |
| *Brain/Cognition* | 1314 | 96 | 0.131 |
| *Ear* | 134 | 10 | 0.332 |
| *Endocrine Metabolic* | 452 | 25 | 0.937 |
| *Eye* | 347 | 17 | 0.924 |
| *Face* | 252 | 29 | 0.004* |
| *GI tract* | 159 | 11 | 0.441 |
| *Genitalia* | 67 | 8 | 0.062 |
| *Heart Cardiovasculature/Lymphatic* | 240 | 32 | 2.62E-04* |
| *Kidney/Renal Tract* | 148 | 11 | 0.511 |
| *Musculature* | 124 | 11 | 0.162 |
| *Respiratory tract* | 33 | 2 | 0.291 |
| *Skeleton* | 687 | 47 | 0.577 |
| *Skin* | 241 | 23 | 0.054 |
| *Spinal cord/Peripheral nerves* | 57 | 2 | 0.764 |
| *Teeth and Dentition* | 50 | 0 | 0.966 |
| *All OMIM* | 3433 | 259 |  |

To follow up on these results, we investigate whether the LIMBR scores can be used in a case-control study to improve our ability to detect gene-disease associations. In practice, the scores could potentially be used as a weighting factor in order to increase power in disease related gene detection. Using a cohort of 488 epileptic encephalopathy cases and 12,151 unrelated controls from a previous rare variant collapsing analysis^23^, we performed a group-wise association test on the five epilepsy genes mentioned above (Eq 4).^24^ In the first set of tests, we assign equal weights to all variants; in the second, we weight the variants by the inverse percentile of their exon-level intolerance scores (details in Methods). Four out of the five genes appear to demonstrate lower p-values under the weighting scheme, while the weighted p-value for KCNT1 increases but has overlapping 95% confidence intervals with the unweighted p-value (Figure 7f). This serves as a proof of concept for the potential to use the scores to aid in identifying key genes associated with diseases.

## Discussion

We demonstrate the approach to intolerance scoring implement in LIMBR results in a significant improvement over existing methods in the prediction of pathogenic variants. We believe that the primary reason for this improvement is that LIMBR is able to capture patterns of variation in intolerance amongst genic regions while minimizing the noise introduce by sub-region size. Of particular relevance to diagnostic sequencing LIMBR also identifies a set of 259 genes in which there is a significant relationship between intolerance scores and the location of pathogenic missense mutations. This work therefore identifies a set of existing genes for which diagnostic labs should not only consider location in interpreting variants but should also consider the intolerance of the regions within which variants fall. We anticipate that more effective use of the location of missense variants in genes will help to address substantial challenge presented by variants of unknown significance. Interestingly, some of the genes that have high rates of variants of unknown significance show significant regional variation in intolerance including one gene central to a high-profile court case resulting from misclassification of a pathogenic variant as a variant of unknown significance.^29,30^

As with any analytical model, the analysis is only as good as the constraints of the data. As control data sets increase in size, so will the power to detect intolerance. Furthermore, improvement in genic sub-region definitions and the increasing availability of trait-specific data will also improve the ability to understand sub-regional intolerance.

Beyond the immediate contribution to variant interpretation, the implicit sharing of information across lower levels in the hierarchy allows the statistical framework utilized here to be extended to address a number of topics that have not yet been investigated. The regression formulation we propose is flexible and can accommodate additional covariates; for example, one can include additional levels in the hierarchy to model intolerance within and between populations or species. This would result in improved precision via the sharing of information across groups, but also the ability to estimate population specific effects. This creates exciting opportunities for example to develop an intolerance scoring system trained for example on human and mouse population genetic data which would identify genes that are under common and divergent patterns of purifying selection. LIMBR provides a natural framework for such studies which could provide important insights in the use of animal models to study human genetic disease. The model could be adapt to jointly examine missense and LOF variants as different outcomes, or to incorporate prediction of how damaging individual variants are.^31,32^. We do not go into a detailed comparison of variant-level predictors of pathogenicity^11,33,34^ in this paper because many of these methods use information on pathogenic variants to develop their scores, and they also may not account for gene-level effects. These scores may still be of interest in conjunction with LIMBR for variant interpretation, especially by integrating methods that use orthogonal information to characterize how damaging a variant is. We are currently working on expanding our tests to explore these avenues, and as more data becomes available we will be able to directly address regional variation in more specialized settings.

The Bayesian framework also allows for different priors to be put on the model, potentially leading to shrinkage of genes or sub-regions to improve classification of novel variants. Overall, the Bayesian hierarchical intolerance model is a flexible and robust way to model sub-regional intolerance to variation, showing strong classification of previously identified pathogenic variants while demonstrating less susceptibility to variability in estimates due to sub-region size.

## Online Resources

Online Mendelian Inheritance in Man (OMIM): http://www.ncbi.nlm.nih.gov/omim

Genome Aggregation Database (gnomAD): http://gnomad.broadinstitute.org/

ClinVar aggregation of genomic variation: https://www.ncbi.nlm.nih.gov/clinvar/

